# Bone marrow mesenchymal stem cells stimulated by tumor necrosis factor-α can promote the repair of fatty liver cell oxidative stress injury and fatty liver ischemia-reperfusion injury

**DOI:** 10.1101/2021.11.08.467855

**Authors:** Yuying Tan, Jiali Qiu, Weiqi Zhang, Yan Xie, Chiyi Chen, Junjie Li, Jiang Li, Wentao Jiang

## Abstract

Mesenchymal stem cells (MSCs) have great prospects for the treatment of ischemia-reperfusion injury (IRI) after liver transplantation. At this stage, the main factor limiting MSCs in the treatment of fatty liver IRI of the donor liver is the residence time of stem cells at the site of inflammatory injury. This study investigated whether bone marrow mesenchymal stem cells (BMSCs) stimulated by tumor necrosis factor-α (TNF-α) can promote the repair of fatty liver cell oxidative stress injury and fatty liver IRI in rats. The results indicated the BMSCs treatment group stimulated by TNF-α had lower indexes and significantly improved oxidative stress damage in vitro through Transwell chamber co-culture experiment, compared with the control group. In vivo, compared with the PBS group and the BMSCs group, the indexes of the BMSCs treatment group stimulated by TNF-α were reduced, and the degree of tissue damage was significantly reduced. BMSCs can repair fatty liver cell oxidative stress injury and fatty liver IRI, however, BMSCs stimulated by TNF-α can promote the repair of tissues and cells.

## Introduction

Liver transplantation is the only effective way to treat end-stage liver disease. With the increasing maturity of liver transplantation technology and the increasing number of liver transplant patients, the shortage of donor livers has become more prominent ^1^. How to safely use marginal donor liver has become a research hotspot. As a kind of marginal donor liver, fatty liver has short storage time and is more sensitive to IRI, these problems are more likely to lead to postoperative complications such as primary no function (PNF) and early graft dysfunction^2^. With the increase in the national obesity rate and the aging population in my country, the proportion of fatty liver in the donor liver pool in China is also increasing. Finding a repair method to improve the safety and utilization of fatty liver donor liver is what we urgently and urgently need to solve.

In recent years, more and more studies have shown that MSCs have great prospects for the treatment of IRI after liver transplantation^3,4^. At this stage, the main factor limiting MSCs in the treatment of fatty liver IRI of the donor liver is the residence time of stem cells at the site of inflammatory injury. TNF-α is the main cytokine in the process of inflammation, and MSCs stimulated by cytokine can specifically bind to a large number of selectin ligands accumulated on the microvascular intima at the site of IRI, there is a scientific hypothesis that allows more BMSCs to stay in the ischemic part of the damaged liver^5,6^. Therefore, this experiment aims to explore whether BMSCs stimulated by TNF-α can promote the repair of fatty liver cell oxidative stress injury and fatty liver IRI in rats.

## Result

### Preparation of BMSCs and establishment of oxidative stress model of IAR-20 fatty liver cells

There were specific protein expressions on the surface of BMSCs. According to the expression results of CD79, CD45, CD90, and CD29 detected by flow cytometry, the extracted cells were determined to be BMSCs (FIG. 1 A), which can be followed up for research.

**Fig 1.**
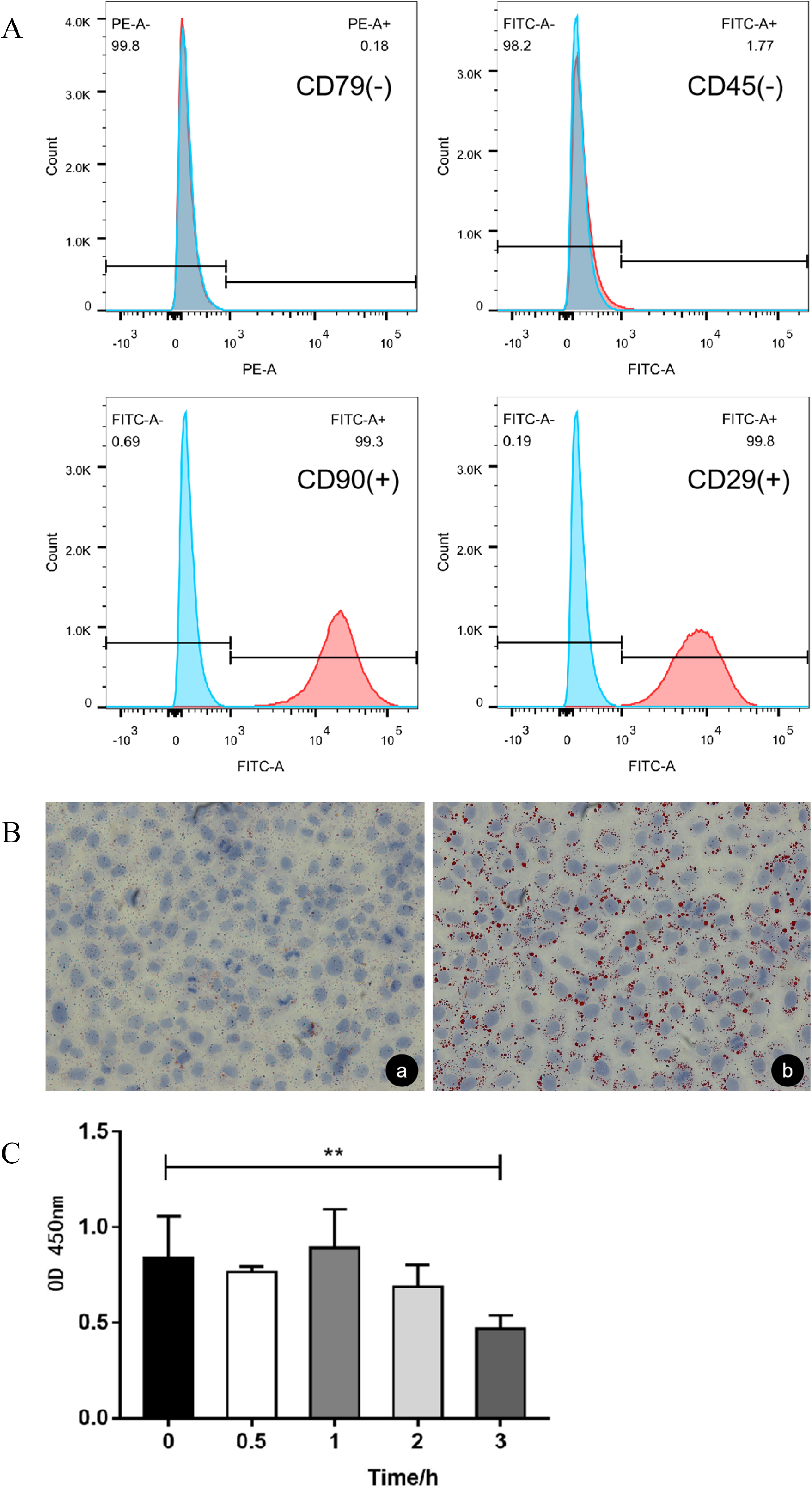
Preparation of BMSCs and optimal concentration for establishing an oxidative stress model. (A)Flow cytometry detection: The expression of CD79, CD45, CD90, and CD29 on the surface of BMSCs(Mesenchymal stem cells have high expression of CD90 and CD29, and low expression of CD79 and CD45).(B)IAR-20 Oil Red O Staining: (a) Normal IAR20 cells are in a small spindle shape, the cytoplasm of liver cells is basically blue, and no red-stained lipid droplets are seen; (b) Fatty IAR-20 cells have a small spindle shape, and a large number of red-stained lipid droplets can be seen in the cytoplasm of liver cells. Original magnification, 100× (C) Cell counting kit-8 assay: Cell viability after H_2_O_2_ stimulates IAR-20. (\*\**P*<0.01)

After stimulating IAR-20 cells with 100μM sodium palmitate and 200μM sodium oleate for 24 hours, a stable fatty liver cell model was formed (FIG. 1 B). After stimulating fatty liver cells with 3mmol/L H_2_O_2_ for 3 hours, the cell viability was significantly reduced, which was about 50% of the cell viability of non-oxidative stress (FIG. 1 C).

These results confirmed that 3mmol/L H_2_O_2_ was the optimal concentration for establishing an oxidative stress model.

### Repair effect of BMSCs stimulated by TNF-α on fatty liver cells damaged by oxidative stress in vitro

After 72 hours of co-cultivation in the Transwell chamber, the supernatant of cells in the lower chamber was collected to detect the function of liver cells and the expression levels of inflammatory factors in each experimental group. Compared with the H-IAR 20 group, the ALT and AST of the T-IAR 20 group and the B-IAR 20 group were significantly lower, but there was no significant difference in the ALB level of each group (FIG. 2 A). Compared with the H-IAR 20 group, the inflammatory indexes of the cell supernatant of the T-IAR 20 group were significantly reduced, and the T-IAR 20 group had a lower level of inflammatory factor expression than the B-IAR 20 group (FIG. 2 B), indicating that compared with the control group, BMSCs stimulated by TNF-α improved the liver function and inflammation of fatty liver cells damaged by oxidative stress.

**Fig 2.**
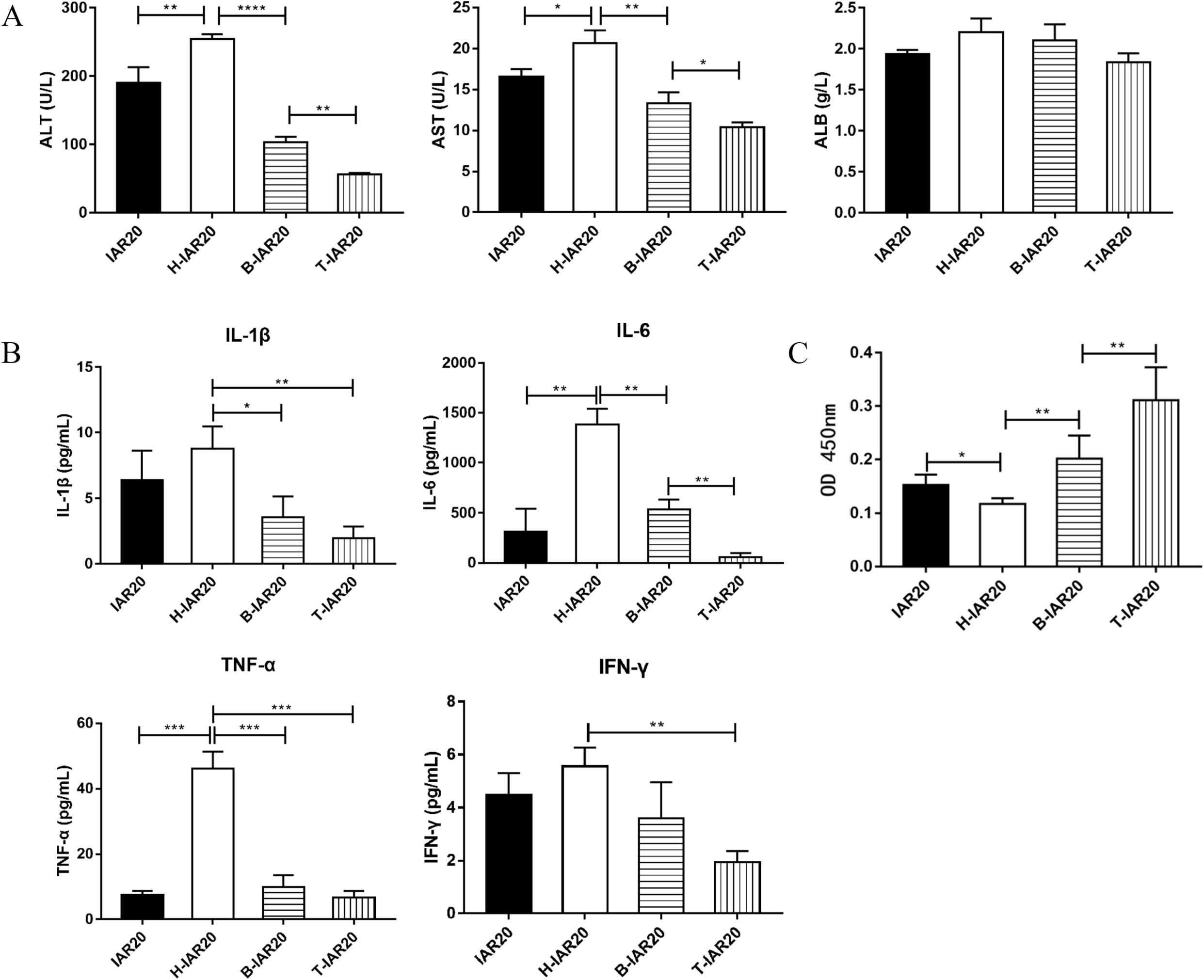
In vitro, effects of BMSCs stimulated by TNF-α on liver function, inflammation and cell viability of fatty liver cells damaged by oxidative stress. (A) Liver function indexes of cell culture supernatant: The liver function indexes ALT and AST of the T-IAR 20 group and the B-IAR 20 group were significantly lower than those of the H-IAR 20 group, and there was no significant difference in the ALB levels of each group. (B) Compared with the H-IAR 20 group, the inflammatory indexes IL-1β, IL-6, TNF-α, and INF-γ of the cell supernatant of the T-IAR 20 group were significantly reduced, and the T-IAR 20 group was compared with the B-IAR 20 group the expression level of inflammatory factors is lower. (C) Cell counting kit-8 assay: Hepatocyte activity of each experimental group(\**P*<0.05, \*\**P*<0.01, \*\*\**P*<0.001, \*\*\*\**P*<0.0001)

CCK-8 assay was used to detect the cell viability of IAR-20, the greater the absorbance, the stronger the cell viability. The results showed that compared with the H-IAR 20 group, the absorbance of the B-IAR 20 group and the T-IAR 20 group was significantly increased, and the cell viability of the injured hepatocytes was significantly enhanced after BMSCs treatment (FIG. 2 C).

Understand cell apoptosis through flow cytometry. According to flow cytometry, the first, second, and fourth quadrants represented cell apoptosis in different periods. Compared with the H-IAR 20 group, the apoptosis rate of hepatocytes in the B-IAR 20 and T-IAR 20 groups was significantly lower (FIG. 3 A, B). The expression levels of HO-1 and bcl-2 represent the anti-oxidative stress and anti-apoptotic ability of cells, respectively. Western blot showed that the expression of HO-1 and bcl-2 proteins in the T-IAR 20 group were significantly increased, and their anti-oxidative stress and anti-apoptotic capabilities were enhanced (FIG. 3 C, D). The above results indicate that in in vitro experiments, BMSCs stimulated by TNF-α have a repairing effect on fatty liver cells damaged by oxidative stress.

**Fig 3.**
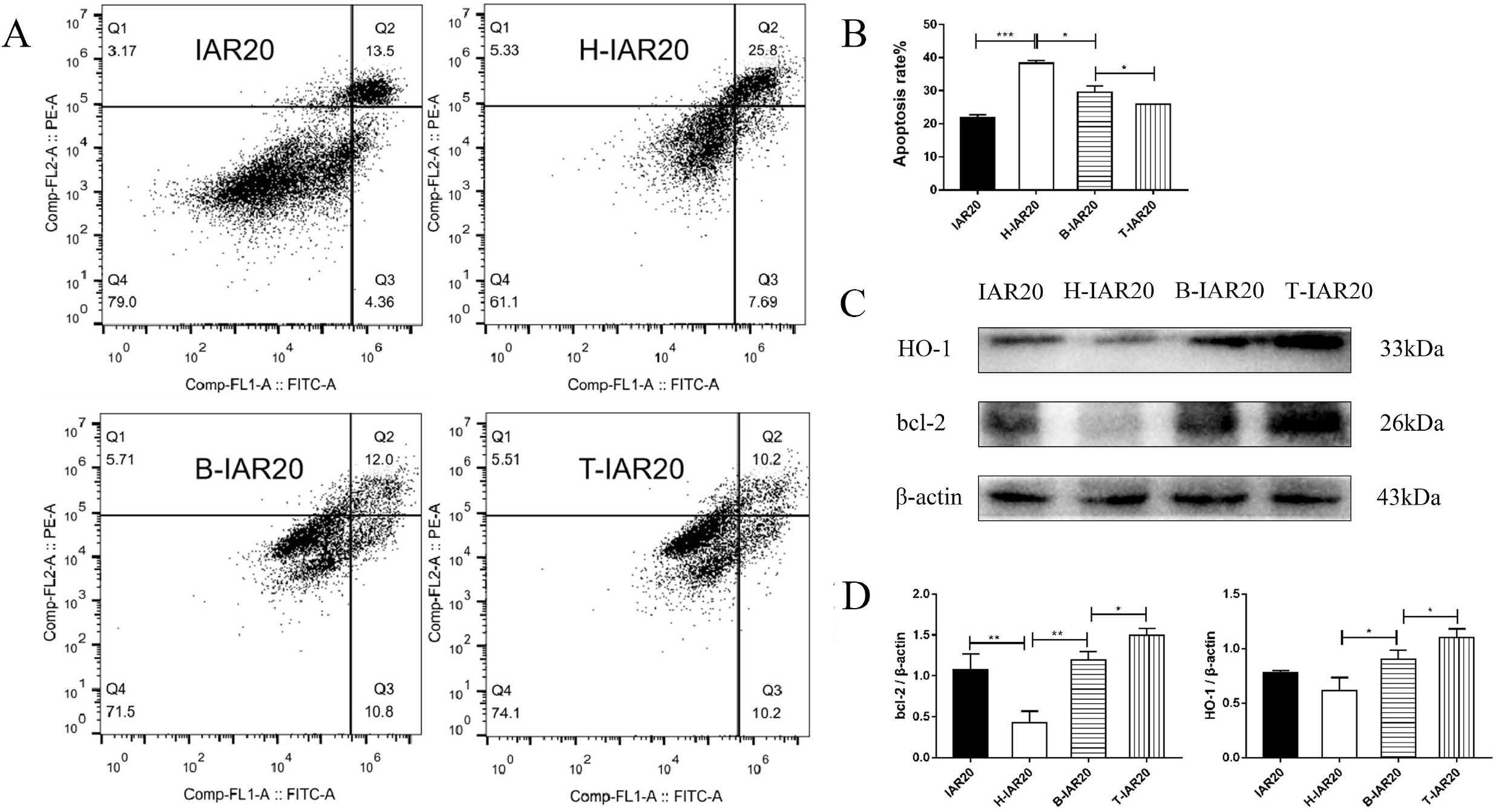
In vitro, effects of BMSCs stimulated by TNF-α on anti-apoptosis and anti-oxidant of fatty liver cells damaged by oxidative stress. (A) Hepatocyte apoptosis of each experimental group. (B) The specific apoptosis rate of hepatocytes in each experimental group. (C) Western blot of HO-1 and bcl-2 in IAR-20, The expressions of HO-1 and bcl-2 were higher in T-IAR 20 group. (D)The bar graph of western blot result for HO-1 and bcl-2.(\**P*<0.05, \*\**P*<0.01, \*\*\**P*<0.001)

### Repair effect of BMSCs stimulated by TNF-α on IRI of fatty liver in rats in vivo

The fatty liver model of rats was established by high-fat diet, and the degree of fatty liver was confirmed by oil red O staining. The proportion of steatosis in liver lobules reached more than 3/4, showing diffuse steatosis, which is severe fatty liver (FIG. 4 A). An ischemia-reperfusion model was artificially constructed by surgery (FIG. 4 B). Finally, it was seen that the clipped liver lobe was dark red and large congestion areas appeared.

**Fig 4.**
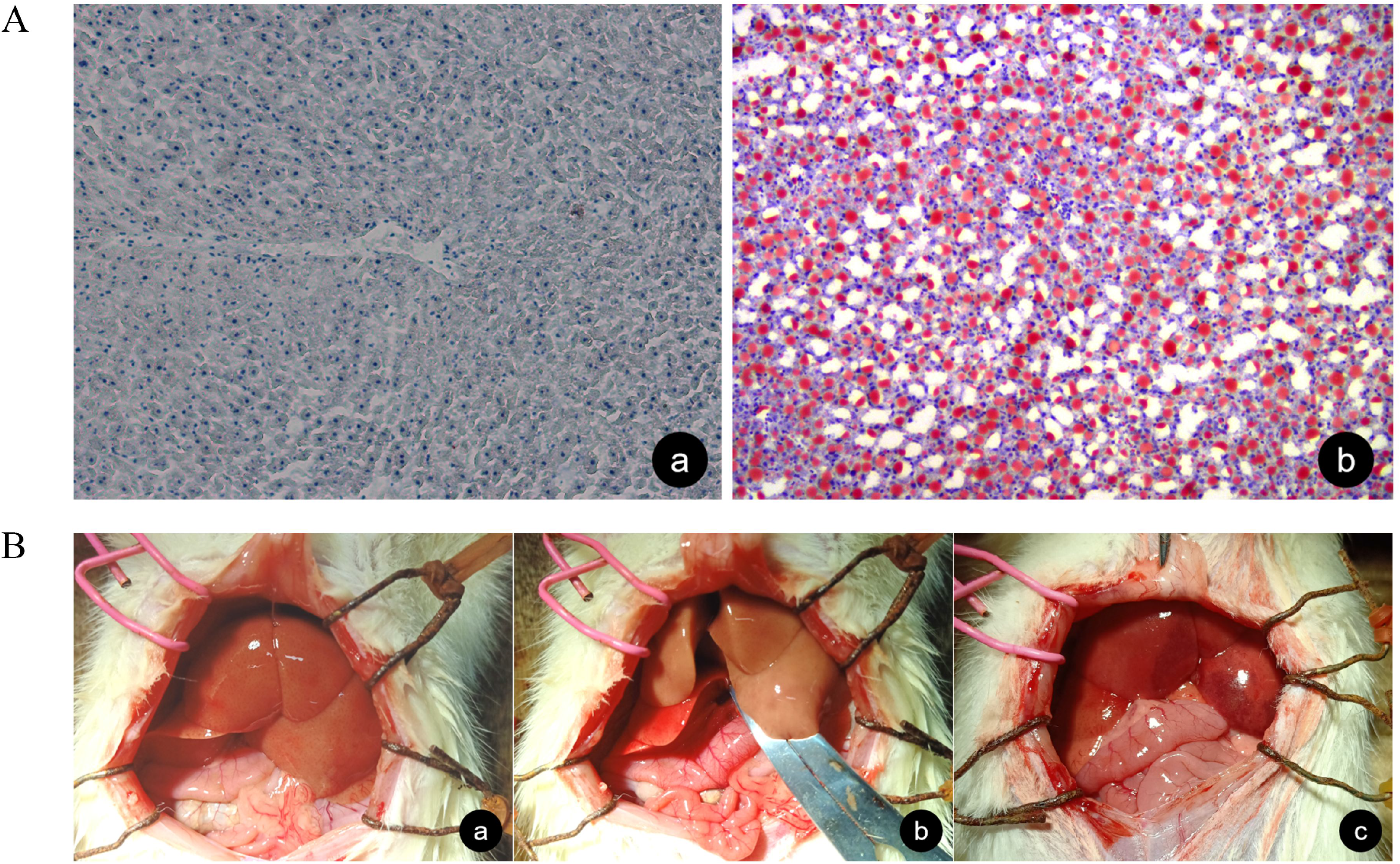
Establishment rat fatty liver and ischemia-reperfusion injury models. (A) Oil Red O staining of liver tissue of SD rats: (a) SD rats fed with normal maintenance feed. (b) SD rats were fed with MCD feed for 2 weeks. (B) SD rat fatty liver ischemia-reperfusion injury before, during and after operation: (a) Before clamping the vascular trunk. (b) Begin to clamp the vascular trunk. (c) Clamp the vascular trunk for 30 minutes and open the blood flow.

12 hours after the treatment of ischemia-reperfusion injury in rats, tissues were collected for detection of liver function, inflammatory factors and other related indexes.The ALT index level of the BMSCs/TNF-α treatment group was significantly lower than that of the PBS treatment group. The TBil index level of the PBS treatment group was significantly higher than that of the Sham group. After treatment with BMSCs, the TBil expression level decreased, while the TBil index of the BMSCs/TNF-α treatment group decreased more significantly, and there was no significant change in AST expression (FIG.5 A). The expression of inflammatory factors in the PBS treatment group was significantly higher than that in the sham group, the BMSCs treatment group and BMSCs/TNF-α treatment group were compared with the PBS treatment group, the inflammatory indexes TNF-α and IL-6 were significantly reduced, however IL-1β and INF-γ was not significantly different (FIG. 5 B). Therefore, the BMSCs/TNF-α treatment group had a greater impact on liver function and its inflammatory factors, which can improve liver function status and reduce inflammation. In rat liver tissues, the expression levels of bcl-2 and HO-1 proteins tended to be consistent with the in vitro experimental results. The expressions of the two proteins in the BMSCs treatment group and the BMSCs/TNF-α treatment group were significantly higher than those in the PBS treatment group (FIG.5 C, D), BMSCs/TNF-α also exerted strong anti-oxidation and anti-apoptosis abilities in vivo.

**Fig 5.**
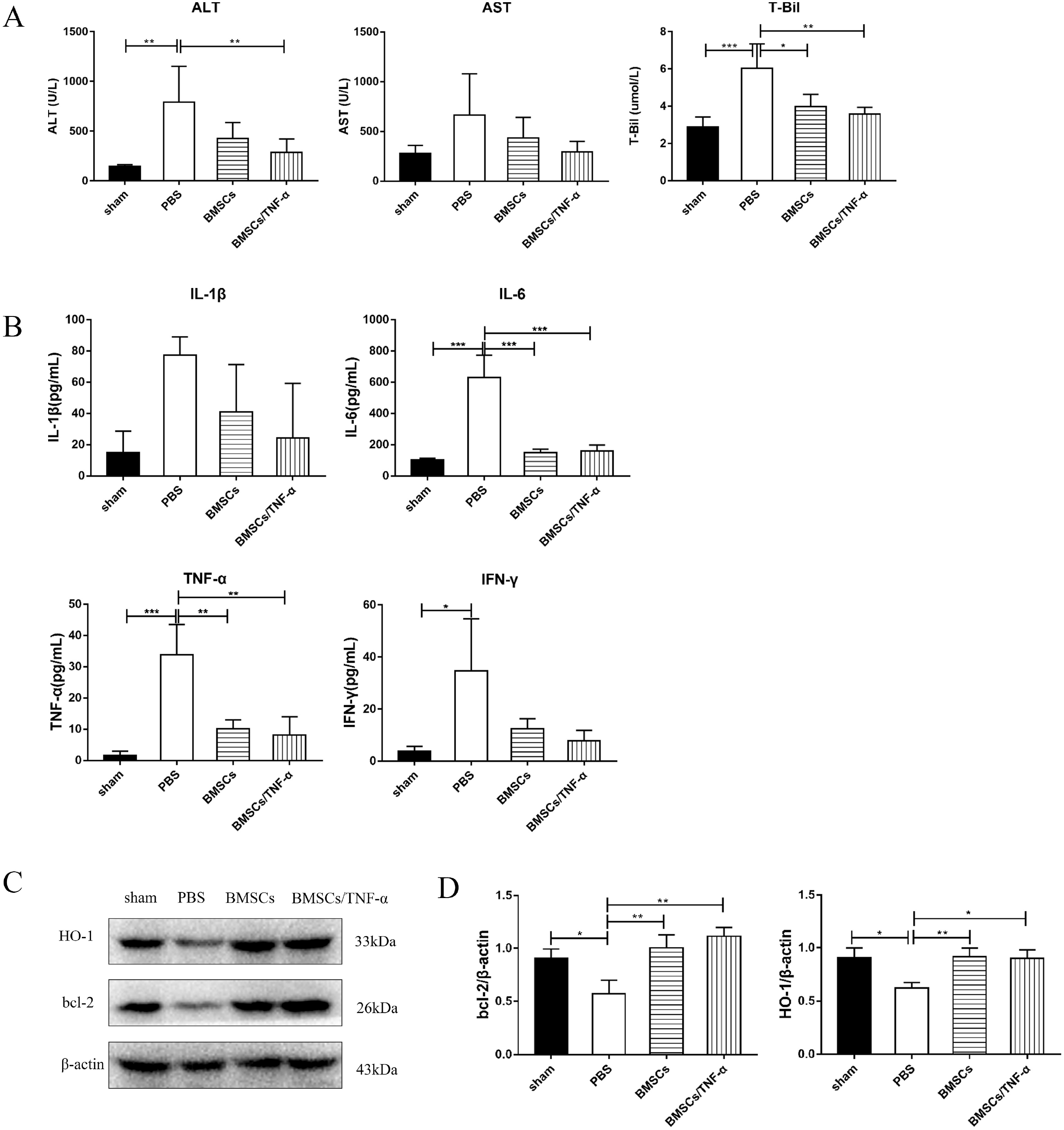
In vivo, the effects of BMSCs stimulated by TNF-α treatment on liver function, inflammation, anti-oxidation and anti-apoptosis of ischemia-reperfusion injury. (A) Serum liver function index levels in SD rats of each experimental group: Compared with the ALT index level of the Sham group, the ALT index of the PBS treatment group was significantly increased, and the BMSCs/TNF-α treatment group was significantly lower than that of the PBS treatment group; P The TBil index level of the PBS treatment group was significantly higher than that of the Sham group. After treatment with BMSCs, the TBil expression level decreased, while the TBil index of the BMSCs/TNF-α treatment group decreased more significantly; There was no significant change in AST expression. (B) Serum inflammation index levels in SD rats of each group. (C) Western blot of HO-1 and bcl-2 in rat liver tissue, the expression of bcl-2 and HO-1 protein in the PBS group was significantly lower than that in the Sham group, the expression of bcl-2 and HO-1 protein in the BMSCs treatment group and the BMSCs/TNF-α treatment group was significantly higher than that of the PBS treatment group. However, the differences in the expression of their own proteins in the first two groups were not significant. (D) The bar graph of western blot result for HO-1 and bcl-2.(\**P*<0.05, \*\**P*<0.01, \*\*\**P*<0.001)

The results of pathological HE staining of liver tissues in each group showed that the BMSCs/TNF-α treatment group reduced liver tissue damage to a greater extent than the other groups (FIG. 6 A). Injecting CM-Dil labeled BMSCs into SD rats can more intuitively observe the status of BMSCs in the liver. It was observed by fluorescence microscope that the number of BMSCs in the liver in the BMSCs/TNF-α treatment group was significantly higher than that in the BMSCs treatment group (FIG. 6 B, C).

**Fig 6.**
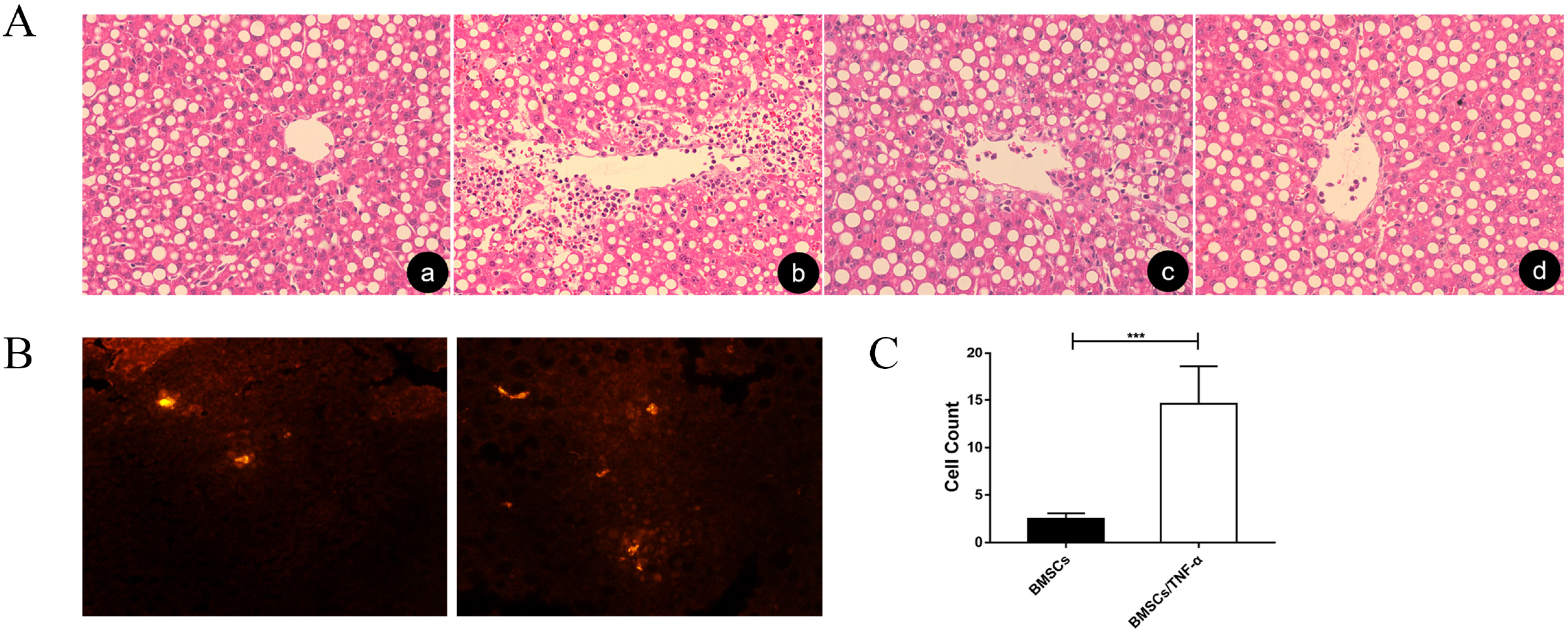
The effect of different treatment methods on the liver tissue of IRI and the retention of BMSCs in the liver of rats. (A) HE staining of liver tissue: (a)In the Sham group, there was a large number of balloon-like changes and a small amount of inflammatory cell infiltration, showing the appearance of moderate to severe fatty liver; (b) In the PBS treatment group, loss of liver structure, hepatic cord disintegration, hemorrhage, hepatocyte necrosis and disintegration, nuclear pyknosis or even disappearance, neutrophil infiltration, and a large number of inflammatory cell infiltration; (c) In the BMSCs treatment group, the boundaries of liver cells were blurred, the structure of liver lobules was disordered, the hepatic sinusoids were congested, and there were more inflammatory cell infiltrations. Compared with the PBS group, the inflammatory response was reduced and cell necrosis was restored; (d) The BMSCs/TNF-α treatment group showed slight damage to the liver lobule structure, mild local congestion, and inflammatory cell infiltration. The degree of liver cell damage, congestion, and inflammation were alleviated compared with the PBS treatment group and the BMSCs treatment group. (B) Fluorescence coloration of BMSCs in liver tissue (a BMSCs treatment group; b BMSCs/TNF-α treatment group). (C) The number of BMSCs cells per high-power field of view. Original magnification, 200×(\*\*\**P*<0.001)

These results showed that, whether in vivo or in vitro, TNF-α can enhance the functions of cell homing and repair by stimulating BMSCs, and promote the anti-inflammatory, anti-oxidant and other repair effects of IRI.

## Discussion and Conclusion

MSCs are widely distributed in various parts of the body and are a type of adult stem cells with self-renewal and multi-differentiation potential. It can participate in the repair process and participate in the inflammatory response ^7^ through mechanisms such as cell fusion or differentiation, paracrine^8^, gene carriers, immunomodulatory function^9^, homing and initiation of endogenous repair. BMSCs are mesenchymal stem cells that exist in the bone marrow stroma without hematopoietic function. Compared with other MSCs, BMSCs have unique advantages such as relatively easy extraction and weak immunogenicity^10^, it can proliferate indefinitely in vitro and has the potential to differentiate into tissue cells derived from mesoderm and neuroectoderm.

BMSC repairs oxidative stress damage through a variety of mechanisms: such as secretion of growth factors, regulation of cell and tissue activity through cytokines and chemokines, and immune regulation ^12–15^. Related in vitro experimental studies have shown that co-culture of hepatocytes and BMSCs can better maintain the original shape of hepatocytes, and maintain the ability of hepatocytes to synthesize albumin and carry out ammonia metabolism at a higher level^16^, studies have also shown that co-culture of liver cells and MSCs can enhance liver cell function and maintain liver cell metabolism ^18^, CAMUSSI G ^19^and others believe that the soluble factors produced by BMSCs, the secreted extracellular matrix components, and the ratio of hepatocytes/BMSCs co-culture are all involved in the protection of hepatocytes, in vitro experiments using porcine hepatocytes showed that when hepatocytes/BMSCs were co-cultured at a ratio of 2:1, hepatocytes had the best ability to synthesize albumin and urea. Yu-Ting He ^20^ and others used hepatocytes/MSCs at a culture ratio of 5:1 for culture to generate stable primary hepatocytes. MINHUI LI ^21^and others co-cultured hUc-MSC and HK-2 cells at a ratio of 12:1 to prove that hepatocyte growth factor derived from human umbilical cord blood mesenchymal stem cells promoted autophagy in HK-2 cells treated with AOPP. Studies have considered that the co-culture system can make BMSCs differentiate into hepatocytes to a certain extent ^21,25^, but the co-culture experiment of BMSCs such as Lange has proved that the proportion of BMSCs differentiation into hepatocytes is very small in a relatively short period of time^26^, the impact on the experiment is not large. Therefore, in this experiment, the indirect co-cultivation method was used to co-culture IAR-20 and BMSCs, and the co-cultivation ratio was 5:1.

The BMSCs stimulated by cytokines can produce specific protein receptors, which bind to the large number of selectin ligands accumulated on the microvascular intima of the injured site, so that more BMSCs can act on the injured site, and further repair the tissue. The stimulated MSCs are further activated to further activate the functions of paracrine and immune regulation, and promote tissue repair. It has been reported in the literature that TNF-ɑ can promote the proliferation and migration of BMSCs ^26^, he ^27^ and others proved that low concentrations of IFN-γ can promote the proliferation and migration of dental pulp MSCs, and inhibit their differentiation. Koning ^28^and others proved that inflammatory factors play an important role in stimulating the migration of MSCs. Essid ^29^ and others found that TNF-ɑ at a concentration of 20 ng/ml can induce rat hepatocyte apoptosis. In this experiment, 10 ng/ml TNF-ɑ was selected to pre-stimulate BMSCs, which can improve the proliferation and migration ability of BMSCs to a certain extent, further activate BMSCs, and prevent inflammatory factors from inducing rat hepatocyte apoptosis.

The liver IRI during liver transplantation includes two parts: cold and warm IRI30. In this experiment, a surgical model established by simulating liver warm IRI was to clamp part of the blood supply vessels of the rat liver to achieve 70% of the liver tissue ischemia^31^, and open the blood flow after 30 minutes of clamping. Using this method to establish a model can achieve effective liver tissue damage and ensure the survival rate of rats during surgery.

MSCs have been proved to be safe and effective for clinical treatment, but the treatment effect of MSCs alone is weak. MSCs died in the first few hours of transplantation, and their homing efficiency was low due to in vitro culture and other reasons. Therefore, improving the survival rate and homing ability of MSCs in vivo is an important part of improving the therapeutic effect of MSCs. Methods such as overexpression of chemokines^32^ and increasing the expression level of adhesion molecules^33^ are used to explore and improve the homing, survival, and function of BMSCs. In this experiment, the inflammatory factor TNF-α was used to pre-stimulate BMSCs for 24 hours to promote the expression of cell surface-related adhesion molecules and activate cell functions, so as to enhance the homing ability of BMSCs and the anti-inflammatory and anti-oxidant repair effects on damaged liver tissues.

In summary, after low-dose TNF-α pre-stimulated BMSCs, the ability of BMSCs to repair the oxidative stress damage of fatty liver cells in vitro had been significantly enhanced, and TNF-α enhanced the functions of cell homing and repair by stimulating BMSCs, and promoted the anti-inflammatory and anti-oxidant repair effects of HIRI. Therefore, this experiment provides theoretical basis and new ideas for optimizing the clinical treatment plan, improving the therapeutic effect of stem cells, and repairing liver IRI.

## Materials and Methods

### Experimental animals

Healthy SPF male SD rats, 8-10 weeks old, weighing 200-250g, and healthy SPF male SD rats, 2-4 weeks old, weighing 40-60g, purchased from China Food and Drug Control Research Institute, the culture environment did not exceed 5 per cage, free drinking water, the breeding environment temperature was (24±2) °C, and the relative humidity was (60±10) %.

### Ethics approval

All animal experiments were approved by the Experimental Animal Ethics Committee of Nankai University and processed according to the national and international rules of animal welfare.

### The preparation, identification and pretreatment of BMSCs

Male SD rats of 2 to 4 weeks old and weighing 40 to 60 g were sacrificed under anesthesia with sodium pentobarbital (Hubei Hongyunlong Biotechnology Co., Ltd.) and placed in 75% alcohol for 15 minutes, under aseptic conditions, separated the femur and tibia, cutted off the epiphyses on both sides, used a 5ml syringe to flush out the cells, filtered twice with a 40μm sieve, added 5ml of red blood cell lysate, placed at 4°C for 10-15 minutes, stopped the lysis and centrifuged to collect the buffy coat cells, resuspended in MEM-α medium (HyClone, USA) containing 15% FBS (HyClone, USA) and antibiotics (100 U/mL penicillin G and 100 mg/mL streptomycin), cultivated in a 37°C, 5% CO2 cell incubator, and cultivated to the third generation for use.

Selected the 3rd generation of well-growing BMSCs to adjust the cell concentration to 5×10^5^ cells/100μl, and added anti-CD 29-PE, anti-CD 90-FITC, anti-CD 45-FITC, and anti-CD 79-PE (eBioscience, USA), incubated for 30 min at room temperature in the dark, washed with PBS (HyClone, USA), centrifuged, and resuspended in 100μl PBS. Used flow cytometry to detect the expression levels of the four proteins on the surface of BMSCs cells.

BMSCs were cultured to the third generation, 10ng/ml TNF-α (Peprotech, USA) was added to the cell culture medium of BMSCs, and cultured in a cell incubator for 24 hours.

### Cell culture and establishment of oxidative stress injury model of fatty liver cells

Mouse normal liver cells IAR-20 were purchased from the Cell Resource Center of the Institute of Basic Medicine, Chinese Academy of Medical Sciences. Cultured in MEM/EBSS (HyClone, USA) medium containing 10% FBS and antibiotics (100 U/mL penicillin G and 100 mg/mL streptomycin) in a 37°C, 5% CO2 cell incubator.

Inoculated normal IAR-20 in a 96-well plate, the number of cells was about 1×10^4^ cells/well, used high-fat complete medium containing 10% FBS and antibiotics, 200μM sodium oleate (Sigma O-7501, USA)-100μM sodium palmitate (Sigma P-9767, USA) for 24h. After the hepatocytes were fatty, 3mmol/L H2O2 was added for stimulation, the stimulation time gradient was set to 0h, 0.5h, 1h, 2h, 3h, and the best stimulation time was selected.

### Co-cultivation of Transwell Chamber System and Grouping of Cell Experiments

Used transwell chambers (corning, USA) for indirect co-cultivation. The upper chamber was BMSCs (2×10^5^/well) and the lower chamber was IAR-20 (1×10^6^/well). The total culture was 72 hours, and each group had 3 replicate wells, taked out the cell, collected the cell culture supernatant, washed twice with PBS, trypsin (Solarbio, China) digestion and centrifugationed to collect the hepatocytes in each well. According to different co-cultivation systems, the cells were divided into fatty IAR-20 group (IAR 20 group), oxidative stress damaged fatty IAR-20 group (H-IAR 20 group), oxidative stress damaged fatty IAR-20 and BMSCs co-culture group (B-IAR 20 group), fatty IAR-20 damaged by oxidative stress and BMSCs co-culture group (T-IAR 20 group) stimulated by TNF-α in 4 groups.

### Establishment of fatty liver IRI model and grouping of animal experiments

Choose 8-10 weeks old male SD rats weighing 200-250g, give Methionone- and Choline-deficient (MCD) model feed (Nantong Trofe Feed Technology Co., Ltd.) diet for 2 weeks to establish a stable fatty liver model. Intraperitoneal injection of 1% sodium pentobarbital (40mg/kg) to maintain the anesthesia of the donor rat, clamped the left outer and left middle hepatic artery trunks, and opened the blood vessels after clamping for 30 minutes, used an insulin needle to inject 0.5ml of the treatment liquid through the portal vein, close the abdominal cavity, and put it into the postoperative recovery box for resuscitation. Grouped according to different treatment methods: ① Sham operation group (Sham group),②Fatty liver IRI PBS treatment group (PBS group),③BMSCs treatment group for fatty liver IRI (BMSCs group),④BMSCs stimulated by TNF-α treatment group for fatty liver IRI(TNF-α/BMSCs group).

### Detection index

Liver function indexes: ALT, AST, ALB, using ELISA kit (Youda, China) to detect inflammation indicators: IL-1β, IL-6, TNF-α, INF-γ, the luminescence situation of CM-Dil in BMSCs.

### CCK-8 assay

After 72 hours of co-cultivation, the cells from the lower chamber were collected and added to a 96-well plate. The number of cells was about 5000 cells/well. Added 10μl of CCK-8 (Solarbio, China) to each well, incubated for 1h in a cell incubator, and detected the OD value at a wavelength of 450nm using a microplate reader.

### ANNEXIN V-FITC/PI detects cell apoptosis

After 72 hours of co-cultivation, collected the IAR-20 cells in the six-well plate, washed with PBS, suspended the cells with 1ml of 1× Binding Buffer, centrifuged and discarded the supernatant, and then resuspended the cells with 1× Binding Buffer. Adjusted the cell density to 1×10^6^ cells/ml, took 100μL of cell suspension, added 5μL Annexin V-FITC (Solarbio, China) to the tube, at room temperature, protected from light, mixed gently, 10min, then added 5μL PI, at room temperature, protected from light, and incubated for 5 min, added PBS to 500μL, mixed gently, and detected cell apoptosis by flow cytometry within 1 hour.

### Western blotting

Electrophoresis was performed using 10% SDS-PAGE (Solarbio, China) and transferred to PVDF membrane (Millipore, USA). Next, the membrane was sealed in 5% skimmed milk powder (BD, USA) for 2h, Antibodies used for immunoblotting in this study were specific to HO-1, bcl-2 (Abcam, USA) and β-actin protein antibody (Proteintech, USA). Used enhanced chemiluminescence detection system to detect target protein.

### Statistics

GraphPad 8.0.2 statistical software was used to analyze and graph the data. The measurement data was expressed as mean ± standard deviation. The independent Student’s t test was used to compare the differences between the two groups, analysis of variance compares the differences between two or more groups, A P value <0.05 means significant difference, and a P value <0.01 means extremely significant difference. Each experiment was repeated 3 times.

## Author contributions

Funding acquisition, Wentao Jiang; Investigation, Yuying Tan and Jiali Qiu; Methodology, Weiqi Zhang, Yan Xie and Jiang Li; Project administration, Wentao Jiang; Visualization, Chiyi Chen and Junjie Li; Writing – original draft, Yuying Tan, Jiali Qiu and Weiqi Zhang; Writing – review & editing, Wentao Jiang.

